# Characteristic of KPC-14, a KPC Variant Conferring Resistance to Ceftazidime-Avibactam in the Extensively Drug-resistant ST463 *Pseudomonas aeruginosa* Clinical Isolate

**DOI:** 10.1101/2024.11.14.623663

**Authors:** Yanyan Xiao, Le Wang, Huiqiong Jia, Yan Jiang, Yue Li, Jiamin Han, Shengchao Li, Yaxi Gu, Qing Yang

## Abstract

We studied two KPC-14 variants from clinical *Pseudomonas aeruginosa* isolates, C137 and C159, to better understand genomic diversity, mechanisms, and genes that confer antibiotic resistance and pathogenicity. C137 and C159, a sequence type 463 ExoU-positive multidrug-resistant strain, were concurrently resistant to carbapenems and ceftazidime-avibactam (CZA). Both strains possessed five intrinsic antimicrobial resistance genes (*fosA, catB7, crpP, bla*_PAO_, and a *bla*_OXA-486_ variant), as well as the *bla*_KPC-14_ gene in the chromosome. In strain C137, *bla*_KPC-14_ gene was situated on the plasmid pC137. KPC-14-harbouring transformants of pC137 exhibited resistance to CZA and restored sensitivity to carbapenems, signifying a “see-saw” effect. Both strains demonstrated the expression of the *bla*_KPC-14_ gene, concurrent inactivation of the outer membrane protein OprD, overexpression of the efflux pump MexX, and a pronounced capacity for biofilm formation. The genomic environment of KPC-14 consisted of IS*26*/IS*26*/TnpR_*Tn3*/IS*Kpn27*/IS*Kpn6*/IS*26*, which was classified as pseudo-compound transposons (PCTs). Plasmid pC137 closely resembled the previously described plasmid p94, which possesses the same genomic architecture, implying that IS*26*-mediated PCTs may store a variety of resistance genes, including *bla*_KPC-2_ and KPC variants, and were currently disseminating in the region. The KPC-14 variant presents significant challenges for clinical treatment. The *bla*_KPC-14_ gene, carried by the PCTs, was integrated into the chromosome, exhibiting stability throughout bacterial inheritance. Our research highlights the necessity for improved clinical surveillance of *Klebsiella pneumoniae* carbapenemase-producing *P. aeruginosa*.

---

Carbapenemase-producing *Pseudomonas aeruginosa* (CP-PA) has caused a worldwide epidemic that is ongoing and intensifying. The predominant carbapenemase types identified in *P. aeruginosa* worldwide are Ambler Class B metallo-enzymes, including VIM, IMP, and NDM^[1]^. The frequency of CP-PA has significantly increased in China, according to recent studies^[2]^, especially in the East China region, where the KPC-2 subtype is most commonly detected^[3]^ and the ST463 type has become the most common clonal type of CP-PA in China^[4]^.

Ceftazidime-avibactam (CZA), including KPC-producing strains, displayed notable effectiveness against carbapenem-resistant *P. aeruginosa* ^[5]^. Nevertheless, since the widespread clinical use of CZA, KPC-producing strains develop resistance to CZA through mutation and the presence of various other factors, and the KPC variant emerged as the main resistance mechanism^[6, 7]^. 216 KPC variants have been identified in the NCBI database(https://www.ncbi.nlm.nih.gov/pathogens/refgene/#*bla*KPC). KPC-31^[8]^, KPC-33^[9]^, KPC-90^[10]^, and KPC-113^[11]^ are the KPC variants of *P. aeruginosa* that have been clinically reported thus far. Numerous KPC variations have been found in in vitro induction studies using CZA^[12, 13]^. The appearance of KPC variants after CZA therapy is concerning and calls for continued observation. It has been noted that *P. aeruginosa* acquires a variety of mobile genetic elements (MGEs) by horizontal gene transfer. Among these elements are unit transposons, integrative and mobilizable elements (IMEs), integrative and conjugative elements (ICEs), and derivatives of IMEs^[11, 14-16]^, these components are essential to the spread of resistance genes.

The insertion sequence IS*26* exhibits significant activity in propagating antibiotic-resistance genes. IS*26* can integrate a gene or a collection of genes into a mobile gene pool and facilitate its dissemination to new sites by producing pseudo-compound transposons (PCTs). The research conducted by Christopher et al. delineated multiple mechanisms by which IS*26* integrates into chromosomes or plasmids: untargeted copy-in cointegration, targeted conservative cointegration, and neighboring deletion or inversion^[17]^.

Limited reports have been issued on KPC variants in *P. aeruginosa*. The epidemiology, resistance mechanisms and pathogenicity of KPC variations in *P. aeruginosa* remain insufficiently studied. In this work, we used genomic and molecular genetic approaches to show that KPC-14, a two-amino-acid deletion in loop 237-24, mediated CZA resistance in CP-PA and identified the mechanisms that contributed to antimicrobial resistance.

## MATERIALS AND METHODS

### Strain isolation and antimicrobial susceptibility testing

In 2022, two *P. aeruginosa* strains, C137 and C159, demonstrating significant resistance to CZA, were isolated from a tertiary hospital in Hangzhou, China. Antimicrobial susceptibility testing was conducted via broth microdilution methods. The results were interpreted according to the breakpoints recommended by the European Committee on Antimicrobial Susceptibility Testing and the Clinical and Laboratory Standards Institute (CLSI)^[18, 19]^.

### Whole-genome sequencing

Genomic DNA of C137/159 were subjected to short-read sequencing (Illumina, San Diego, California, USA) and MinION long-read sequencing (Oxford Nanopore Technologies, Oxford, United Kingdom). Hybrid de novo genome assembly was executed with Unicycler v0.5.0, followed by annotation using Bakta and Prokka 1.14.6^[20]^ and subsequent manual review via BLASTn/BLASTp^[21]^. Multilocus sequence typing (MLST), resistance, and virulence genes were identified using Resfinder^[22]^ and ABRicate v1.0.1 (https://github.com/tseemann/abricate). Transposons were characterized using the transposon registry^[23]^, ISfinder^[24]^, and INTEGRALL^[25]^. Plasmid typing was achieved with PlasmidFinder2.1 (https://cge.food.dtu.dk/services/PlasmidFinder/). Circular genome maps and comparative genomic maps were generated with Easyfig and Brig^[26]^. The origin of mobile genetic element DNA sequence transfer was determined using OriTfinder (https://toolmml.sjtu.edu.cn/oriTfnder/oriTfnder.html). Phylogenetic analysis was performed using CSI Phylogeny (https://cge.food.dtu.dk/services/CSIPhylogeny/) and BacWGSTdb (http://bacdb.cn/BacWGSTdb/).

### Cloning experiments

Recombinant expression vectors were produced utilizing molecular cloning methods. The *bla*_KPC-14_ gene, together with its upstream promoter region, was amplified from pC137, while the previously sequenced *bla*_KPC-2_ and *bla*_KPC-33_ genes served as references for amplification. Their sequences were cloned into the previously documented pBAD24 plasmid and subsequently transformed into *Escherichia coli* DH5α for the assessment of minimum inhibitory concentrations (MICs).

### Checkerboard assay

To elucidate the function of avibactam (Avi) synthesized by KPC-14-producing strains in conferring resistance to CZA, various doses of Avi (0, 4, 8, 16 μg/mL) were combined with a gradient of ceftazidime (CAZ) concentrations (1 to 1024 μg/mL) via a series of two-fold dilutions. The combination was assessed utilizing a checkerboard assay in accordance with CLSI M07 standards, concentrating on strains C137/159 and KPC-14-harboring transformants of pC137. *P. aeruginosa* strains carrying KPC-2/33 and its transformants functioned as reference strains, whereas *P. aeruginosa* ATCC27853 acted as the quality control strain.

### Detection of carbapenemase activity

NG-test Carba 5 was applied for carbapenemase detection in strains C137/159 and KPC-14-harboring transformants of pC137. The modified carbapenem inactivation method (mCIM) followed the CLSI M100 guidelines.

### Gene transcription analysis by reverse transcription-quantitative PCR

The total RNA from bacterial cells was extracted using the Steadypure Universal RNA Extraction Kit to evaluate the expression of efflux pumps (MexAB-OprM, MexCD-OprJ, MexEF-OprN, and MexXY-OprM), the AmpC enzyme, and the outer membrane protein OprD. The RNA was reverse transcribed into cDNA following the kit’s protocol. Subsequently, the relative mRNA expression levels of these genes were determined using a Bio-Rad real-time PCR system. The primers used are listed in Table S1, and reactions were repeated in triplicate. An increase in expression was indicated by a relative expression level >2 for *mexA* and >5 for *mexC, mexE*, and *mexX*^[27]^. For *ampC* and *oprD*, an increase was noted when the relative expression level exceeded 2^[28]^.

### Efflux pump inhibition

The MICs of CZA were determined in the presence and absence of PA*β*N (Phe-Arg-*β*-naphthylamide) (Sigma Aldrich) at a concentration of 50 mg/L^[29]^. The isolates were confirmed to overexpress efflux pumps when the MICs in the presence of PA*β*N were determined to be at least 4-fold lower than the MICs in the absence of PA*β*N^[30]^. The wild-type *P. aeruginosa* PAO1 strain was used as the reference strain.

### Growth curves and biofilm formation assay

The same amount (1×10^6^ CFU/mL) of mid-logarithmic phase ancestral or evolved strains were cultured in 200 μL fresh LB in 96-well plates^[31]^. The plates were incubated in BioTek Synergy H1 Multi-Mode Microplate Reader at 37°C with continuous shaking for 24 h. OD_600_ nm was measured every 15 min. All experiments were carried out three times. *P. aeruginosa* ATCC27853 was used as control, and *P. aeruginosa* strains carrying KPC-2 (P2034) and KPC-33 (P2079) were utilized as reference strains. Growth curves were plotted using OriginPro 2024b, and maximum growth rates were tested using R scripts. As described previously, a biofilm formation test was performed on a Microtiter 96-well plate^[32]^. All experiments were carried out thrice, and the solution’s optical density (OD) at 570 nm was measured. The ability of biofilm formation was classified as weak (ODs ⩽ 2ODc) or strong (ODs > 2ODc). [ODc is the mean OD of the negative control; ODs is the mean OD of samples].

## RESULTS

### Clinical Background

From the sputum samples of two lung transplant patients (A and B Patient), C137 and C159 were identified. After the transplant, Patient A had a lengthy hospital stay. Because of changes in his clinical status, CZA had to be given to him more than once during this time. Patient B and Patient A were hospitalized in the same intensive care unit over the same period, although Patient B had no history of administering CZA after the transplant.

### Antimicrobial susceptibility and genome sequencing analysis

Both C137 and C159 were exclusively susceptible to amikacin and colistin. Resistance was observed against all tested β-lactams and β-lactamase inhibitors, including carbapenems, CZA, imipenem/relebactam, and meropenem/vaborbactam, classifying them as XDR *P. aeruginosa* (Table 1).

**TABLE 1.**
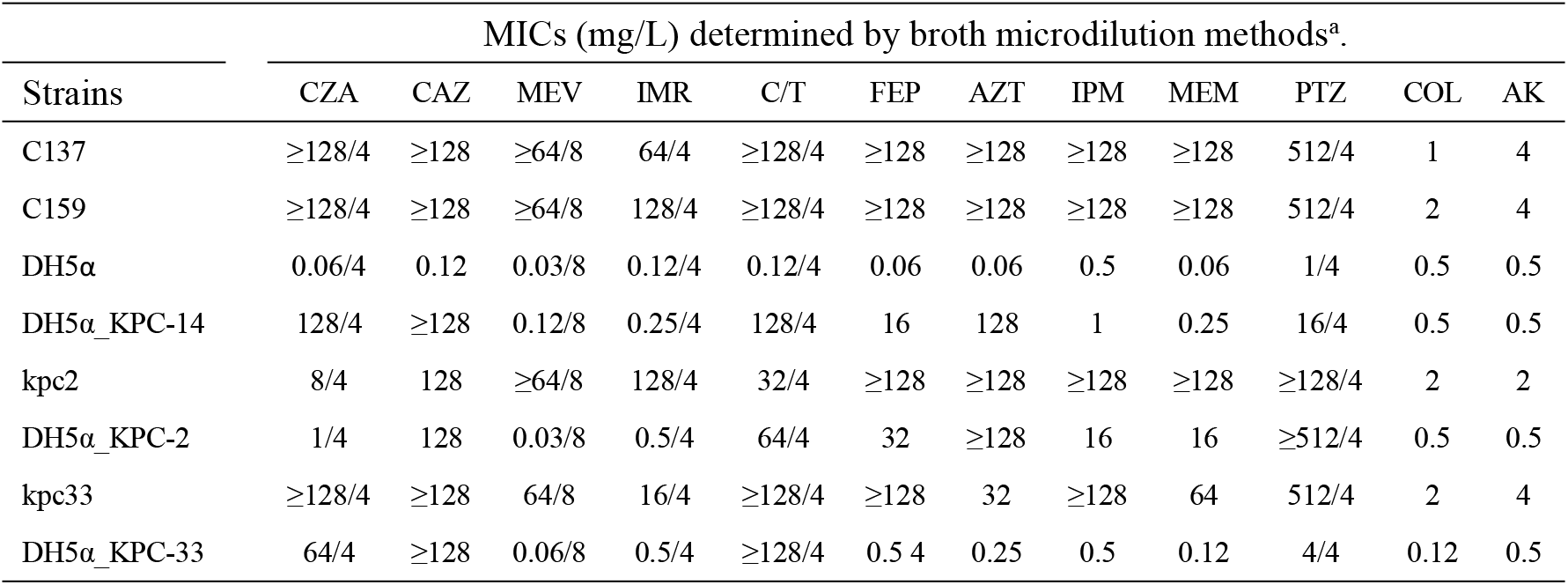
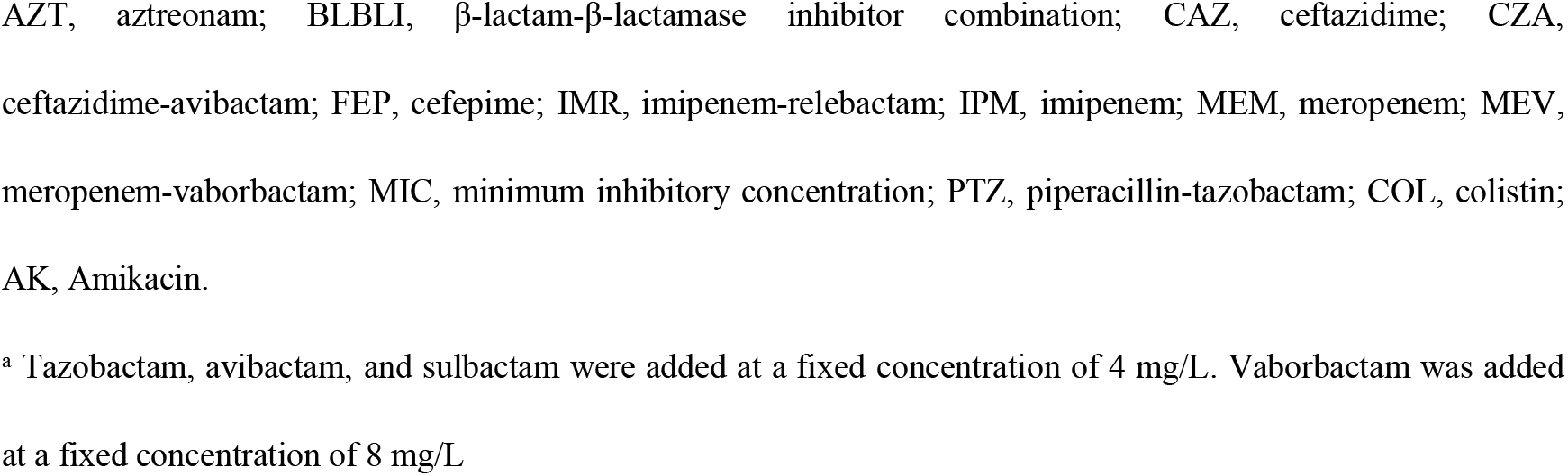
MICs of β-lactams and BLBLIs for strains used in this study.

| Strains | MICs (mg/L) determined by broth microdilution methods<sup>a</sup>. |  |  |  |  |  |  |  |  |  |  |  |
| --- | --- | --- | --- | --- | --- | --- | --- | --- | --- | --- | --- | --- |
|  | CZA | CAZ | MEV | IMR | C/T | FEP | AZT | IPM | MEM | PTZ | COL | AK |
| C137 | $\geq 128/4$ | $\geq 128$ | $\geq 64/8$ | 64/4 | $\geq 128/4$ | $\geq 128$ | $\geq 128$ | $\geq 128$ | $\geq 128$ | 512/4 | 1 | 4 |
| C159 | $\geq 128/4$ | $\geq 128$ | $\geq 64/8$ | 128/4 | $\geq 128/4$ | $\geq 128$ | $\geq 128$ | $\geq 128$ | $\geq 128$ | 512/4 | 2 | 4 |
| DH5α | 0.06/4 | 0.12 | 0.03/8 | 0.12/4 | 0.12/4 | 0.06 | 0.06 | 0.5 | 0.06 | 1/4 | 0.5 | 0.5 |
| DH5α_KPC-14 | 128/4 | $\geq 128$ | 0.12/8 | 0.25/4 | 128/4 | 16 | 128 | 1 | 0.25 | 16/4 | 0.5 | 0.5 |
| kpc2 | 8/4 | 128 | $\geq 64/8$ | 128/4 | 32/4 | $\geq 128$ | $\geq 128$ | $\geq 128$ | $\geq 128$ | $\geq 128/4$ | 2 | 2 |
| DH5α_KPC-2 | 1/4 | 128 | 0.03/8 | 0.5/4 | 64/4 | 32 | $\geq 128$ | 16 | 16 | $\geq 512/4$ | 0.5 | 0.5 |
| kpc33 | $\geq 128/4$ | $\geq 128$ | 64/8 | 16/4 | $\geq 128/4$ | $\geq 128$ | 32 | $\geq 128$ | 64 | 512/4 | 2 | 4 |
| DH5α_KPC-33 | 64/4 | $\geq 128$ | 0.06/8 | 0.5/4 | $\geq 128/4$ | 0.5 4 | 0.25 | 0.5 | 0.12 | 4/4 | 0.12 | 0.5 |

AZT, aztreonam; BLBLI, β-lactam- β-lactamase inhibitor combination; CAZ, ceftazidime; CZA, ceftazidime-avibactam; FEP, cefepime; IMR, imipenem-relebactam; IPM, imipenem; MEM, meropenem; MEV, meropenem-vaborbactam; MIC, minimum inhibitory concentration; PTZ, piperacillin-tazobactam; COL, colistin; AK, Amikacin.
<sup>a</sup> Tazobactam, avibactam, and sulbactam were added at a fixed concentration of 4 mg/L. Vaborbactam was added at a fixed concentration of 8 mg/L

Whole-genome sequencing revealed that the *bla*_KPC-14_ gene was present on the chromosomes of both strains C137 and C159. Furthermore, plasmid pC137 was discovered to contain *bla*_KPC-14_ gene. The chromosomes and plasmid of pC137 were found to share the same KPC genetic contexts (IS*26*/IS*26*/TnpR*_*Tn*3*/IS*Kpn27*/IS*Kpn6*/IS*26*) (Fig.1,2,3). Chromosomes harboured five intrinsic antimicrobial resistance genes (*fosA, catB7, crpP, bla*_PAO_, and a *bla*_OXA-486_ variant). The *bla*_OXA-486_ gene was integrated into the chromosome by IS*26*, while no other resistance genes were associated with genetic elements. Only the *bla*_KPC-14_ gene was present on the plasmid pC137 (37,679 bp), which was devoid of any virulence or antibiotic resistance genes.

**FIG 1.**
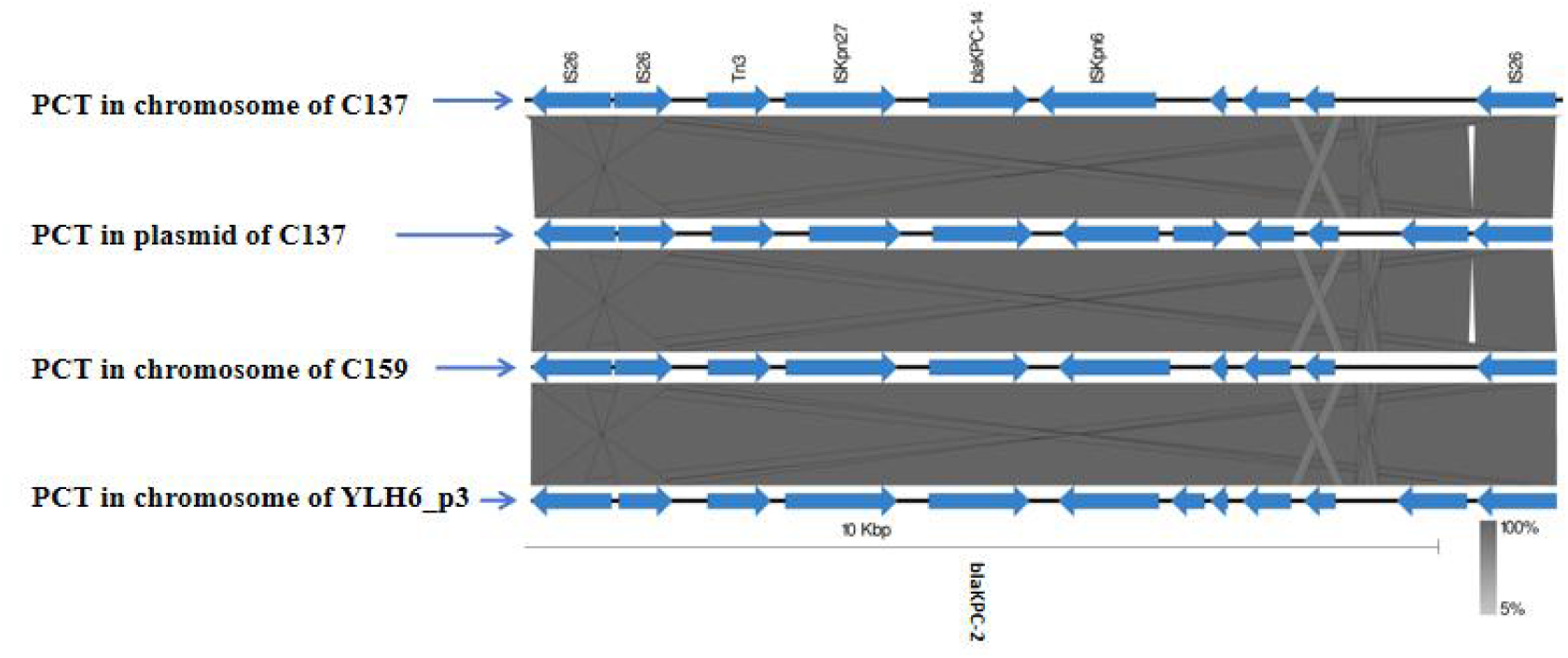
A comparative structural map of the PCT elements found in the chromosomes and plasmids of C137/159 and YLH6_p3. The PCT structures are characterized by IS*26*/IS*26*/TnpR_*Tn3*/IS*Kpn27*/*bla*_KPC-14_/2/IS*Kpn6*/IS*26*.

**FIG 2.**
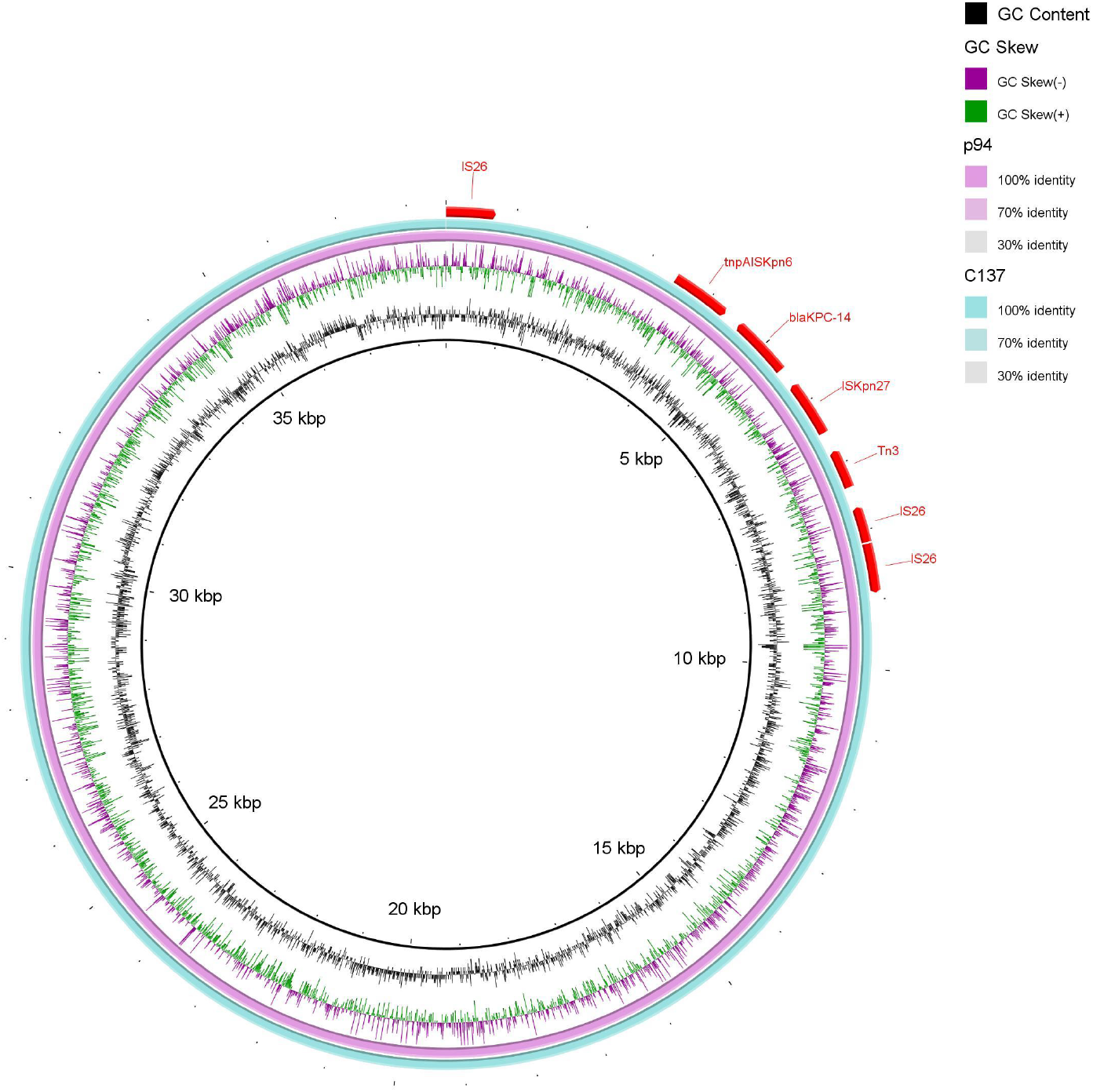
Genetic organization of the pC137 and comparison with the p94(Accession No. CP125362) plasmid.

**FIG 3.**
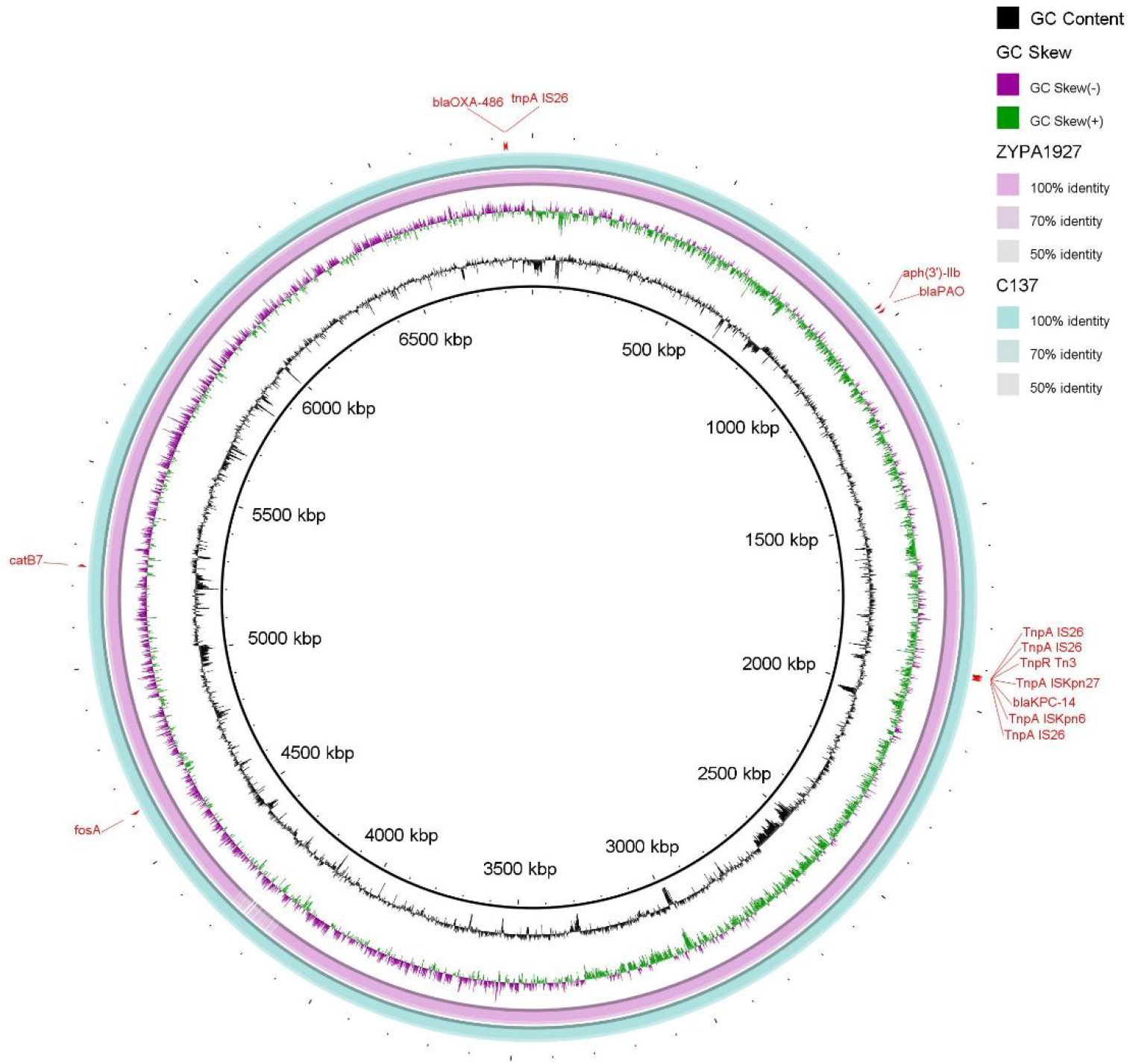
Genetic organization of the chromosome of C137 and comparison with the chromosome of ZYPA1927 (Accession No. CP133751.1).

PlasmidFinder was unable to classify the plasmid containing *bla*_KPC-14_ gene in pC137, which was identified as a non-conjugative plasmid. Analysis performed with oriTfinder showed that the C137 plasmid lacked DNA transfer initiation and relaxation enzymes. The antimicrobial susceptibility tests of the transformants showed sensitivity to imipenem (IPM, MIC 0.25 μg/mL) and meropenem (MEM, MIC 0.12 μg/mL). Nonetheless, the MIC values for ceftazidime (CAZ) and CZA continued to exhibit resistance relative to the C137/159 parental strain, while the MIC of cefepime was reduced from ≥128 μg/mL to 16 μg/mL. The checkerboard assay results show that at Avi concentrations of 0, 4, 8, and 16 μg/mL, the CZA concentration for the C137/159 parental strains varied from 512 to ≥1024 μg/mL. In the absence of Avi (0 μg/mL), the transformants demonstrated a CAZ MIC of ≥1024 μg/mL. The transformants exhibited a CZA MIC of 128 μg/mL at an Avi concentration of 4 μg/mL and at Avi dosages of 8 and 16 μg/mL, the CZA minimum inhibitory concentration (MIC) was documented as ≤1 μg/mL.

The PCTs of the plasmid and chromosomal DNA in this investigation were defined by typical 820 bp IS*26* elements at both termini, encompassing tnpR transposase and 14 bp terminal inverted repeats (TIR). An inverted IS*26* copy was discovered subsequent to the left-side IS*26*, indicating a copy-in cointegration and the genetic factor enabling the transfer of KPC-14 discovered in our work is categorized as a PCT. The fundamental structure, Tn*3*-IS*Kpn27*-*bla*_KPC_-IS*Kpn6*, was classified within the Tn*6296* group and the IS*26* facilitates the transfer of transposons containing resistance genes (IS*26*-Tn*6296*-IS*26*).

This research identified both strains as sequence type 463 (ST 463). The strain examined in our study possessed a type III secretion system that produced exoU, exoS, exoT, and exoY, which are now key indications for evaluating *P. aeruginosa* pathogenicity. The exotoxin gene ExoA was also identified. Furthermore, genes associated with pili and flagella, which facilitate adhesion and biofilm formation, as well as genes involved in anti-phagocytosis and immunological evasion, were identified. The isolates possessed numerous virulence factors responsible for iron acquisition, protease synthesis, and quorum sensing (QS).

### Carbapenemase activity

To assess the detectability of various methodologies for KPC-14, the CARBA5 fast test, and the mCIM assay were performed. The CARBA5 techniques demonstrated positive findings for all isolates containing KPC-2/33/14, including the transformants. The mCIM assay indicated carbapenemase positive for all isolates harboring KPC-2/33/14. The transformants harboring KPC-2 tested positive for carbapenemase using mCIM, but those transformants carrying KPC-33/14 yielded negative results for the carbapenemase assay.

### Detection of efflux pump and analysis of OprD (Porin D) protein

All carbapenem drugs were resistant to the parental strain C137/159. Then, the inquiry was expanded to include a look at the possible mechanisms. The *oprD* gene of the C137/159 strain has 126 nucleotide alterations, according to a BLASTn comparison with the *P. aeruginosa* PAO1 strain. Six of these modifications were deletions of nucleotides, eight were insertions, and the remaining alterations were substitutions. The inactivation of OprD5 was revealed by PorinPredict analysis of the outer membrane porins OprD from the two strains (table S2). *P. aeruginosa* PAO1 and C137’s OprD5 protein were compared structurally by superimposition using the Swiss model and the RCSB PDB (https://www.rcsb.org/) tools (Fig. 4). Quantitative real-time PCR results revealed that the expression of the efflux pump gene *mexX* rose, whereas that of the *oprD* gene decreased.

**FIG 4.**
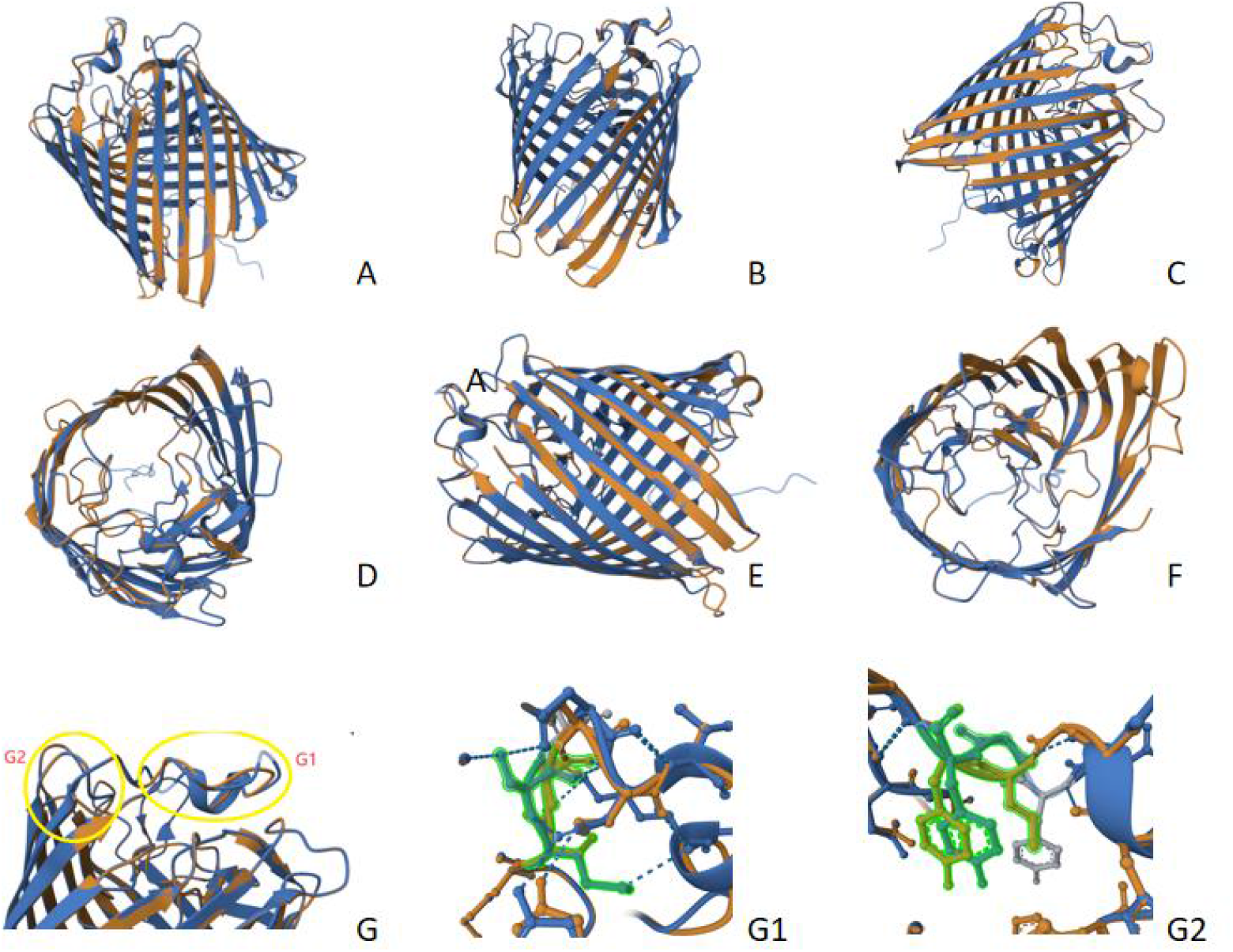
(A-F) illustrates the tertiary structures of outer membrane protein D from *P. aeruginosa* C137 (blue) and PA01 (orange). Panels (G/G1/G2) juxtapose the protein D structures from *P. aeruginosa* C137 and PA01 utilizing the Swiss Model and the RCSB PDB online database. Panels G1 and G2 demonstrate structural modifications at two sites resulting from amino acid substitutions and deletions.

The efflux pump inhibition assays were further performed. We found that CZA susceptibility in the C137/159 isolate was decreased by ≥64-fold in the presence of PA*β*N compared with the absence of Pa*β*N (⩾1024 μg/mL versus 16 μg/mL).

### Growth curves and bioinformatics analysis

*P. aeruginosa* ATCC27853 grew at a greater maximum rate than both the parental strain C137/C159 and the strain harboring KPC-2/33, suggesting that C137/C159’s growth was inhibited in the absence of antimicrobial stress (Fig. 5). The C137/C159 strains were found to be strong biofilm formers.

**FIG 5.**
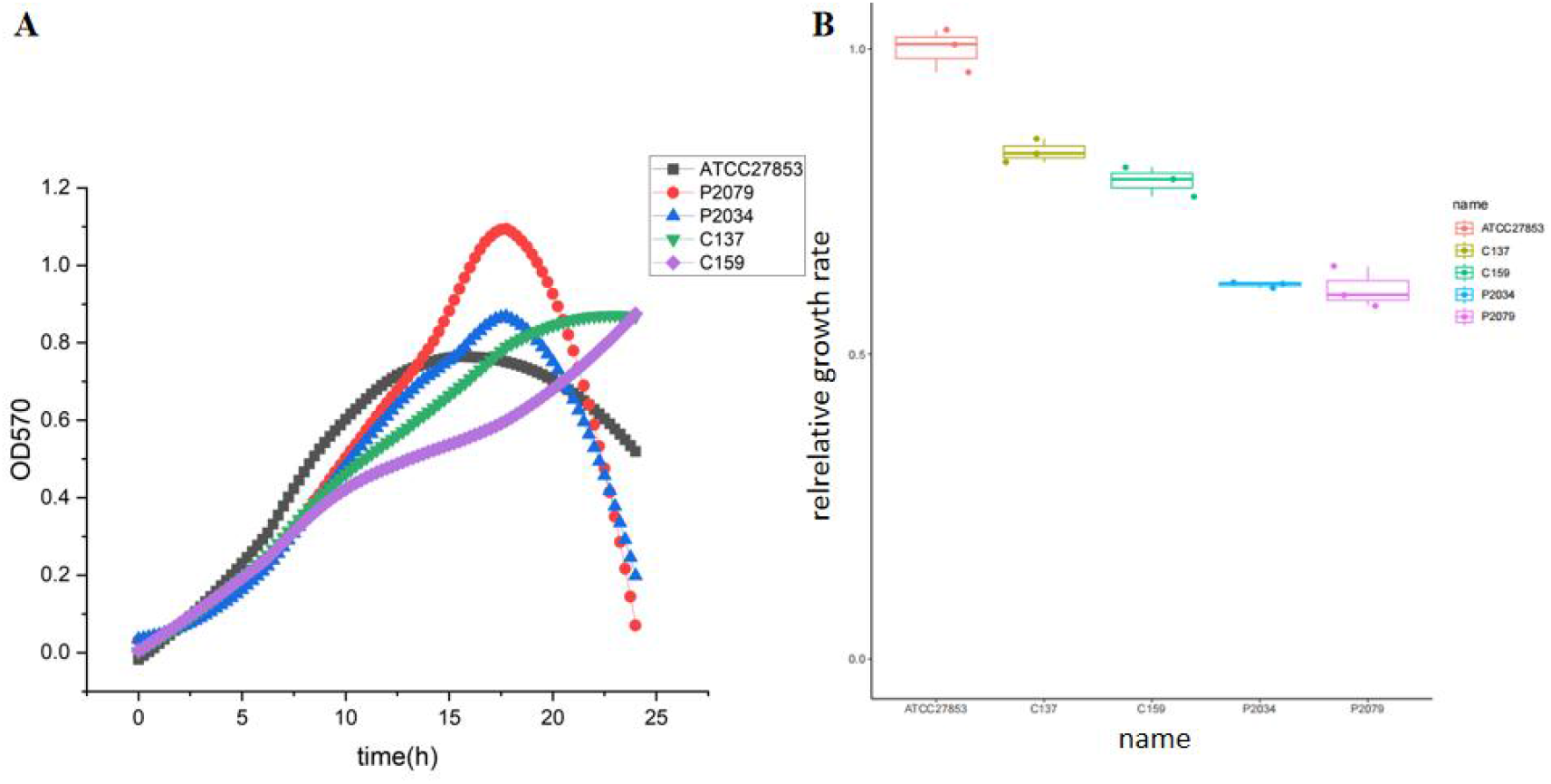
(A) A growth kinetics of C137, C159, P2034, P2079 and ATCC27853; (B) Relative growth rate of C137, C159, P2034, P2079 and ATCC27853 in LB broth medium without antibiotics.

## DISCUSSION

There are currently reports of a significantly higher prevalence of carbapenem-resistant *P. aeruginosa* (CR-PA) in several different parts of the world. Clusters of ST463 ExoU-positive *P. aeruginosa* were found in Southeast China^[14, 33, 34]^, which were linked to a high death rate. A multicenter investigation conducted in China from 2016 to 2019 found that 374 CR-PA strains were isolated from seven hospitals. Of them, 40.4% of the strains produced KPC-2, and 70.9% of the KPC-2 producers were ST463 strains^[2]^. Additionally, in *P. aeruginosa*, lower susceptibility to CZA has been linked to the overexpression of β-lactam enzymes such as KPC-2.

C137/159’s phylogenetic research revealed that they were both descended from the same strain. Both patients were admitted to the ICU at the same time, according to a review of their medical histories, which raises the possibility of a horizontal transfer of the strain while they were receiving treatment.

Results of CZA, IPM, and MEM antimicrobial susceptibility in KPC-14-producing transformants matched those for *Klebsiella pneumoniae* bearing KPC-14^[13, 35-38]^. Similar to what Li et al.^[39]^ found that the susceptibility values for transformants containing KPC-33 were in line with those for *K. pneumoniae* carrying KPC-33. Transformants carrying KPC-14/33 exhibited restored susceptibility to IPM and MEM but retained high-level resistance to CZA. It was discovered that *P. aeruginosa* harboring KPC-14 variant was the source of this phenomenon, which is referred to as the “see-saw” effect. High-level resistance to CZA was first found in studies on several KPC-3 variants (D179Y, T243M, D179Y/T243M, and EL165-166), whereas concurrent restoration of sensitivity to carbapenem was also noted^[40-42]^. The concept of the “see-saw” effect was first discovered in the field with other *K. pneumoniae* KPC variations, including KPC-33. Bacteria interpreted this phenomenon as an evolutionary trade-off in their ability to adapt to antimicrobial stresses.

The majority of KPC variants kept mediating CZA resistance. Three mutation hotspots were identified by Hobson et al. as being associated with CZA resistance: (a) the Ω-loop region (residues 164 to 179), (b) loop 237-243 (located between beta 3 and beta 4), and (c) loop 266-275 (located at a distance from the active site between beta 5 and alpha 11 helix)^[43]^. A key location in KPC is the Ω-loop region, which has many mutations linked to CZA resistance. Further structural changes in the protein may result from mutations at this location, which may improve CAZ’s affinity and accessibility to the active site. This could lead to altered sensitivity to other β-lactams, including carbapenems. Furthermore, certain mutations in this loop can lower the rates of carbamoylation and Avi binding, which can result in resistance to the CZA combination.

KPC-14 was produced by the deletion of two amino acids: glycine at position 242 and threonine at position 243 in KPC-2. The deletions transpired within the range of loop 237 to 243. Oueslati et al. assessed steady-state kinetic characteristics in *E. coli* strains harboring recombinant plasmids for KPC-2/3/14/28. In *E. coli* using recombinant plasmids producing KPC-2/3/14/28, KPC-14 had a heightened affinity for CAZ relative to KPC-2, showing a 40-fold increase in catalytic efficiency. Despite the enhanced affinity for IPM, the catalytic efficiency diminished by 200-fold. These findings suggest that the D242-GT-243 deletion diminishes carbapenem hydrolysis and augments CAZ hydrolysis activity^[44]^.

Cloning experiments indicated that transformants harboring *bla*_KPC-14_ demonstrated increased MIC values for CAZ (≥1024 μg/mL). The augmented resistance of KPC-14 to CZA was chiefly ascribed to the heightened hydrolysis of CAZ, while the specific role of Avi remained ambiguous. The checkerboard experiment demonstrated that for the C137/159 parental strains, the CZA MIC values varied from 512 to ≥1024 μg/mL at all Avi doses but the MIC of CZA for the transformants diminished significantly with elevated Avi concentrations which indicated that the resistance to CZA was mainly attributable to enhanced hydrolysis of CAZ and inadequate concentration of Avi.

The downregulation or inactivation of OprD expression constitutes a mechanism of carbapenem resistance in *P. aeruginosa*^[45]^. OprD, when altered by nonsense or insertion-deletion mutations or disrupted by insertion sequences, leads to IPM resistance^[46]^. The overexpression of efflux pump genes facilitates resistance to multiple antimicrobial drugs in *P. aeruginosa*, including carbapenems^[47]^, and correlates with elevated resistance to CZA^[10, 48]^. Quantitative real-time PCR of our results revealed that *mexX* expression was significantly increased. In the presence of the efflux pump inhibitor, the MIC of the C137/159 isolate was 16 μg/mL, which represented an 8-fold decrease compared to the MIC of the pC137 transformants and an ≥64-fold decrease compared with the absence of Pa*β*N of the C137/159 isolate. This indicates that the efflux pump inhibitor effectively prevented the efflux of Avi, allowing Avi to reach the CAZ binding site and exert its action. These findings further highlight the critical role of the efflux pump in the multidrug resistance mechanisms of *P. aeruginosa* and suggest that insufficient Avi concentration near the CAZ binding site is one of the main contributors to CZA resistance.

The BLASTn findings indicated a 99% query coverage between the chromosomes of C137 and ZYPA1927 (Accession No. CP133751.1), which harbors KPC-2. BLASTn examination of the IS*26*-Tn*6296*-IS*26* area containing KPC-2/14 revealed that the sole distinction between the two was a six-base pair deletion at △ 242-GT-243 in the KPC-14 region. This discovery may indicate that KPC-14 developed from KPC-2, with the PCT harboring *bla*_KPC-2/14_ gene integrating into the bacterial genome. Growth curves demonstrated that both C137 and C159 displayed compromised growth. Following the acquisition of plasmids containing antibiotic resistance, bacterial cells progressively incorporated the resistance genes into their chromosomes to mitigate the fitness costs associated with resistant plasmids^[49]^.

In conclusion, the resistance of KPC-14 to CZA was mostly due to its enhanced hydrolytic activity against CAZ. Moreover, inadequate concentrations of Avi at its target site or its expulsion by efflux pumps likely hindered Avi from successfully accessing the CAZ binding sites, leading to elevated CZA resistance. The inactivation of the outer membrane protein OprD, elevated expression of the efflux pump gene *mexX*, and robust biofilm formation significantly contributed to resistance against carbapenems and CZA. This discovery suggests that the creation of XDR-PA results from the aggregate impact of many resistance mechanisms. Our research demonstrated that PCT harboring KPC-14 was located on plasmids and had integrated into the chromosome, facilitating stable transmission in *P. aeruginosa*. This scenario poses considerable difficulties for the management of multidrug-resistant organisms. Surveillance of local epidemiology might facilitate more effective management of infections caused by XDR-PA.

## ACKNOWLEDGMENTS

This work was supported by the National Natural Science Foundation of China (Grant No. 82073610).

Q.Y. conceived and designed this study. YY.X. and L.W. constructed the plasmids and performed the antimicrobial susceptibility tests. YY.X. and HQ.J. analyzed the genome sequencing data. YY.X. analyzed the modelled protein structure. YY.X. and Y.L. constructed the checkerboard assay. YY.X. JM.H. and YX.G. extracted the RNA and performed the reverse transcription-quantitative PCR. YY.X. and SC.L. performed the growth curves tests and biofilm formation tests. YY.X. and L.W. wrote the initial version of the manuscript. Q.Y., Y.J. and HQ.J. revised the manuscript.

The authors declare that they have no conflicts of interest.

## ETHICS APPROVAL

Ethics approval was obtained from the Ethics Committee of First Affiliated Hospital, Zhejiang University School of Medicine (approval/reference number: IIT20241295A).

## FUNDING

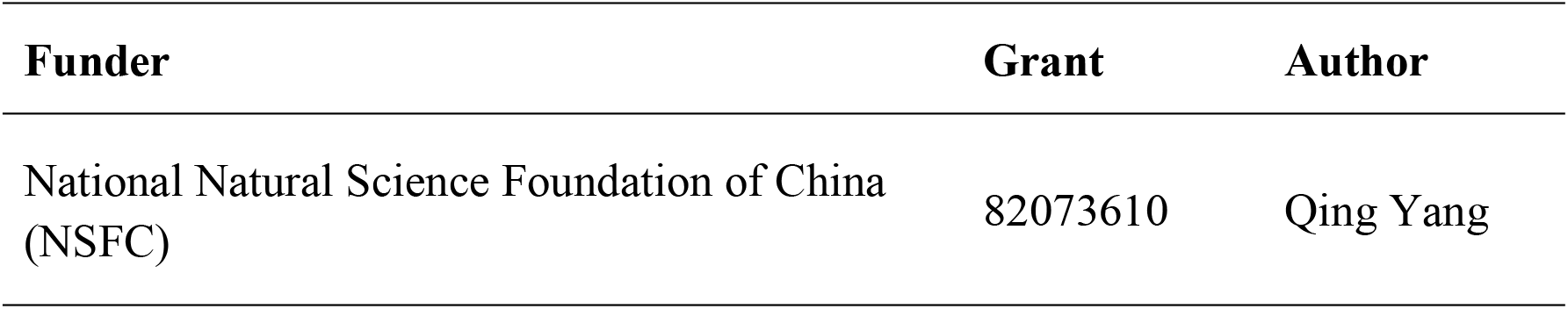

## ADDITIONAL FILES

The following material is available online.

### Supplemental Material

Tables S1 and S2.

## References

[1] Boyd SE, Livermore DM, Hooper DC, et al. Metallo-β-Lactamases: Structure, Function, Epidemiology, Treatment Options, and the Development Pipeline [J]. 2020, 64(10):e00397–20.

[2] Zhu Y, Chen J, Shen H, et al. Emergence of Ceftazidime- and Avibactam-Resistant Klebsiella pneumoniae Carbapenemase-Producing Pseudomonas aeruginosa in China [J]. 2021, 6(6):e0078721.

[3] Yoon EJ, Jeong SH. Mobile Carbapenemase Genes in Pseudomonas aeruginosa [J]. Front Microbiol, 2021, 12:614058.

[4] Hu Y, Liu C, Wang Q, et al. Emergence and Expansion of a Carbapenem-Resistant Pseudomonas aeruginosa Clone Are Associated with Plasmid-Borne bla (KPC-2) and Virulence-Related Genes [J]. 2021, 6(3):e00154–21.

[5] Van Duin D, Bonomo RA. Ceftazidime/Avibactam and Ceftolozane/Tazobactam: Second-generation β-Lactam/β-Lactamase Inhibitor Combinations [J]. Clinical infectious diseases : an official publication of the Infectious Diseases Society of America, 2016, 63(2):234–41.

[6] Ding L, Shen S. Klebsiella pneumoniae carbapenemase variants: the new threat to global public health [J]. 2023, 36(4):e0000823.

[7] Zhang P, Shi Q, Hu H, et al. Emergence of ceftazidime/avibactam resistance in carbapenem-resistant Klebsiella pneumoniae in China [J]. Clin Microbiol Infect, 2020, 26(1):124.e1-.e4.

[8] Faccone D, De Mendieta Juan M, Albornoz E, et al. Emergence of KPC-31, a KPC-3 Variant Associated with Ceftazidime-Avibactam Resistance, in an Extensively Drug-Resistant ST235 Pseudomonas aeruginosa Clinical Isolate [J]. Antimicrobial agents and chemotherapy, 2022, 66(11):e00648–22.

[9] Zhang P, Wang J, Li Y, et al. Emergence of bla(KPC-33)-harboring Hypervirulent ST463 Pseudomonas aeruginosa Causing Fatal Infections in China [J]. The Journal of infection, 2022, 85(4):e86–e8.

[10] Tu Y, Wang D, Zhu Y, et al. Emergence of a KPC-90 Variant that Confers Resistance to Ceftazidime-Avibactam in an ST463 Carbapenem-Resistant Pseudomonas aeruginosa Strain [J]. Microbiology spectrum, 2022, 10(1):e01869–21.

[11] Yang Q, Li Y, Fang L, et al. A novel KPC-113 variant conferring carbapenem and ceftazidime-avibactam resistance in a multidrug-resistant Pseudomonas aeruginosa isolate [J]. Clinical Microbiology and Infection, 2023, 29(3):387.e7-.e14.

[12] Shen S, Tang C, Yang W, et al. In vitro mimicry of in vivo KPC mutations by ceftazidime-avibactam: phenotypes, mechanisms, genetic structure and kinetics of enzymatic hydrolysis [J]. Emerging microbes & infections, 2024, 13(1):2356146.

[13] Jiang M, Sun B, Huang Y, et al. Diversity of Ceftazidime-Avibactam Resistance Mechanism in KPC2-Producing Klebsiella pneumoniae Under Antibiotic Selection Pressure [J]. Infection and drug resistance, 2022, 15:4627–36.

[14] Forero-Hurtado D, Corredor-Rozo ZL, Ruiz-Castellanos JS, et al. Worldwide Dissemination of blaKPC Gene by Novel Mobilization Platforms in Pseudomonas aeruginosa: A Systematic Review [J]. Antibiotics, 2023, 12(4):658.

[15] Fang Y, Wang N, Wu Z, et al. An XDR Pseudomonas aeruginosa ST463 Strain with an IncP-2 Plasmid Containing a Novel Transposon Tn6485f Encoding bla(IMP-45) and bla(AFM-1) and a Second Plasmid with Two Copies of bla(KPC-2) [J]. 2023, 11(1):e0446222.

[16] Yu T, Yang H, Li J, et al. Novel Chromosome-Borne Accessory Genetic Elements Carrying Multiple Antibiotic Resistance Genes in Pseudomonas aeruginosa [J]. Frontiers in cellular and infection microbiology, 2021, 11:638087.

[17] Harmer CJ, Hall RM. IS26 and the IS26 family: versatile resistance gene movers and genome reorganizers [J]. 2024: e0011922.

[18] The European Committee on Antimicrobial Susceptibility Testing. Breakpoint tables for interpretation of MICs and zone diameters. Version 14.0, 2024. [Z].

[19] James S. Lewis Ii P, FIDSA. Performance Standards for Antimicrobial Susceptibility Testing, 34th Edition [Z].

[20] Seemann T. Prokka: rapid prokaryotic genome annotation [J]. Bioinformatics (Oxford, England), 2014, 30(14):2068–9.

[21] Boratyn GM, Camacho C, Cooper PS, et al. BLAST: a more efficient report with usability improvements [J]. Nucleic acids research, 2013, 41(Web Server issue): W29–33.

[22] Zankari E, Hasman H, Cosentino S, et al. Identification of acquired antimicrobial resistance genes [J]. The Journal of antimicrobial chemotherapy, 2012, 67(11):2640–4.

[23] Tansirichaiya S, Rahman MA, Roberts AP. The Transposon Registry [J]. 2019, 10:40.

[24] Siguier P, Perochon J, Lestrade L, et al. ISfinder: the reference centre for bacterial insertion sequences [J]. Nucleic acids research, 2006, 34(Database issue): D32–6.

[25] Moura A, Soares M, Pereira C, et al. INTEGRALL: a database and search engine for integrons, integrases and gene cassettes [J]. Bioinformatics (Oxford, England), 2009, 25(8):1096–8.

[26] Alikhan NF, Petty NK, Ben Zakour NL, et al. BLAST Ring Image Generator (BRIG): simple prokaryote genome comparisons [J]. BMC genomics, 2011, 12:402.

[27] Cabot G, Ocampo-Sosa AA, Tubau F, et al. Overexpression of AmpC and Efflux Pumps in Pseudomonas aeruginosa Isolates from Bloodstream Infections: Prevalence and Impact on Resistance in a Spanish Multicenter Study [J]. Antimicrobial agents and chemotherapy, 2011, 55(5):1906–11.

[28] Tom SM, Doumith M, Warner M, et al. Efflux pumps, OprD porin, AmpC beta-lactamase, and multiresistance in Pseudomonas aeruginosa isolates from cystic fibrosis patients [J]. Antimicrobial agents and chemotherapy, 2010, 54(5):2219–24.

[29] Matsumoto Y, Yamasaki S. Changes in the expression of mexB, mexY, and oprD in clinical Pseudomonas aeruginosa isolates [J]. 2024, 100(1):57–67.

[30] Sonnet P, Izard D, Mulli C. Prevalence of efflux-mediated ciprofloxacin and levofloxacin resistance in recent clinical isolates of Pseudomonas aeruginosa and its reversal by the efflux pump inhibitors 1-(1-naphthylmethyl)-piperazine and phenylalanine-arginine-β-naphthylamide [J]. International journal of antimicrobial agents, 2012, 39(1):77–80.

[31] Xiang G, Zhao Z, Zhang S, et al. Porin deficiency or plasmid copy number increase mediated carbapenem-resistant Escherichia coli resistance evolution [J]. Emerging microbes & infections, 2024, 13(1):2352432.

[32] Zhao J, Zheng B, Xu H, et al. Emergence of a NDM-1-producing ST25 Klebsiella pneumoniae strain causing neonatal sepsis in China [J]. Front Microbiol, 2022, 13:980191.

[33] Zhu Y, Chen J, Shen H, et al. Emergence of Ceftazidime- and Avibactam-Resistant Klebsiella pneumoniae Carbapenemase-Producing Pseudomonas aeruginosa in China [J]. mSystems, 2021, 6(6):e00787–21.

[34] Li Y, Fang L, Dong M, et al. bla (KPC-2) overexpression and bla (GES-5) carriage as major imipenem/relebactam resistance mechanisms in Pseudomonas aeruginosa high-risk clones ST463 and ST235, respectively, in China [J]. 2023, 67(11):e0067523.

[35] Bianco G, Boattini M, Iannaccone M, et al. Carbapenemase detection testing in the era of ceftazidime/avibactam-resistant KPC-producing Enterobacterales: A 2-year experience [J]. Journal of global antimicrobial resistance, 2021, 24:411–4.

[36] Van Asten Sav, Boattini M, Kraakman MEM, et al. Ceftazidime-avibactam resistance and restoration of carbapenem susceptibility in KPC-producing Klebsiella pneumoniae infections: A case series [J]. Journal of infection and chemotherapy : official journal of the Japan Society of Chemotherapy, 2021, 27(5):778–80.

[37] Bianco G, Boattini M, Iannaccone M, et al. Bloodstream infection by two subpopulations of Klebsiella pneumoniae ST1685 carrying KPC-33 or KPC-14 following ceftazidime/avibactam treatment: considerations regarding acquired heteroresistance and choice of carbapenemase detection assay [J]. The Journal of antimicrobial chemotherapy, 2020, 75(10):3075–6.

[38] Niu S, Chavda KD, Wei J, et al. A Ceftazidime-Avibactam-Resistant and Carbapenem-Susceptible Klebsiella pneumoniae Strain Harboring bla(KPC-14) Isolated in New York City [J]. 2020, 5(4):e00775–20.

[39] Li D, Li K. Ceftazidime-Avibactam Resistance in Klebsiella pneumoniae Sequence Type 11 Due to a Mutation in Plasmid-Borne Bla (kpc-2) to Bla (kpc-33), in Henan, China [J]. 2021, 14:1725–31.

[40] Shields RK, Nguyen MH, Press EG, et al. In Vitro Selection of Meropenem Resistance among Ceftazidime-Avibactam-Resistant, Meropenem-Susceptible Klebsiella pneumoniae Isolates with Variant KPC-3 Carbapenemases [J]. Antimicrobial agents and chemotherapy, 2017, 61(5):e00079–17.

[41] Shields RK, Nguyen MH, Press EG, et al. Emergence of Ceftazidime-Avibactam Resistance and Restoration of Carbapenem Susceptibility in Klebsiella pneumoniae Carbapenemase-Producing K pneumoniae: A Case Report and Review of Literature [J]. Open Forum Infectious Diseases, 2017, 4(3):ofx101.

[42] Tsivkovski R, Lomovskaya O. Potency of Vaborbactam Is Less Affected than That of Avibactam in Strains Producing KPC-2 Mutations That Confer Resistance to Ceftazidime-Avibactam [J]. Antimicrobial agents and chemotherapy, 2020, 64(4):e01936–19.

[43] Hobson CA, Pierrat G, Tenaillon O, et al. Klebsiella pneumoniae Carbapenemase Variants Resistant to Ceftazidime-Avibactam: an Evolutionary Overview [J]. 2022, 66(9):e0044722.

[44] Oueslati S, Iorga BI, Tlili L, et al. Unravelling ceftazidime/avibactam resistance of KPC-28, a KPC-2 variant lacking carbapenemase activity [J]. The Journal of antimicrobial chemotherapy, 2019, 74(8):2239–46.

[45] O’Donnell JN, Bidell MR, Lodise TP. Approach to the Treatment of Patients with Serious Multidrug-Resistant Pseudomonas aeruginosa Infections [J]. 2020, 40(9):952–69.

[46] Sherrard LJ, Wee BA, Duplancic C, et al. Emergence and impact of oprD mutations in Pseudomonas aeruginosa strains in cystic fibrosis [J]. Journal of Cystic Fibrosis, 2022, 21(1):e35–e43.

[47] Lorusso AB, Carrara JA, Barroso CDN, et al. Role of Efflux Pumps on Antimicrobial Resistance in Pseudomonas aeruginosa [J]. 2022, 23(24):15779.

[48] Li X, Zhou L, Lei T, et al. Genomic epidemiology and ceftazidime-avibactam high-level resistance mechanisms of Pseudomonas aeruginosa in China from 2010 to 2022 [J]. 2024, 13(1):2324068.

[49] Zongo PD, Cabanel N, Royer G, et al. An antiplasmid system drives antibiotic resistance gene integration in carbapenemase-producing Escherichia coli lineages [J]. Nature Communications, 2024, 15(1):4093.

